# Multi-omics analysis reveals regime shifts in the gastrointestinal ecosystem in chickens following anticoccidial vaccination and *Eimeria tenella* challenge

**DOI:** 10.1101/2024.03.27.586915

**Authors:** Po-Yu Liu, Janie Liaw, Francesca Soutter, José Jaramillo Ortiz, Fiona M. Tomley, Dirk Werling, Ozan Gundogdu, Damer P. Blake, Dong Xia

## Abstract

Coccidiosis, caused by *Eimeria* parasites, poses significant economic and welfare challenges in poultry farming. Beyond its direct impact on health, *Eimeria* infection disrupts enteric microbial populations leading to dysbiosis and increases vulnerability to secondary diseases such as necrotic enteritis, caused by *Clostridium perfringens*. The impact of *Eimeria* infection or anticoccidial vaccination on host gastrointestinal phenotypes and enteric microbiota remains understudied. In this study, the metabolomic profiles and microbiota composition of chicken caecal tissue and contents were evaluated concurrently during a controlled experimental vaccination and challenge trial. Cobb500 broilers were vaccinated with a *Saccharomyces cerevisiae*-vectored anticoccidial vaccine and challenged with 15,000 *Eimeria tenella* oocysts. Assessment of caecal pathology and quantification of parasite load revealed correlations with alterations to caecal microbiota and host metabolome linked to infection and vaccination status. Infection heightened microbiota richness with increases in potentially pathogenic species, while vaccination elevated beneficial *Bifidobacterium*. Using a multi-omics factor analysis (MOFA) machine learning model, data on caecal microbiota and host metabolome were integrated and distinct profiles for healthy, infected, and recovering chickens were identified. Healthy and recovering chickens exhibited higher vitamin B metabolism linked to short-chain fatty acid-producing bacteria, whereas essential amino acid and cell membrane lipid metabolisms were prominent in infected and vaccinated chickens. Notably, vaccinated chickens showed distinct metabolites related to the enrichment of sphingolipids, important components of nerve cells and cell membranes. Our integrated multi-omics model revealed latent biomarkers indicative of vaccination and infection status, offering potential tools for diagnosing infection, monitoring vaccination efficacy, and guiding the development of novel treatments or controls.

## INTRODUCTION

Protozoan parasites of the genus *Eimeria* cause coccidiosis in poultry and costs to the industry have been estimated to exceed £10 billion annually [1]. Clinical coccidiosis manifests as poor body weight gain and feed conversion with diarrhoea, bloody droppings, and mortality in severe cases. Infection induces strong pro- and anti-inflammatory cytokine responses that may exacerbate pathology [2–5]. Clinical coccidiosis is commonly avoided through a combination of good husbandry, parasite chemoprophylaxis with anticoccidial drugs and/or vaccination using varied formulations of live parasites [6, 7]. In some countries, public concern related to pathogen drug resistance and widespread use of antimicrobials in animal production are driving legislative and commercial changes, including increased use of anticoccidial vaccination [8]. Although current live parasite vaccines are effective, considerable efforts are also being made to develop recombinant anticoccidial vaccines [9]. In a previous study, a novel prototype inactivated yeast-based recombinant oral vaccine for *Eimeria tenella* was shown to result in reduced parasite replication, reduced caecal pathology and improved chicken performance compared to controls [10]. Using *Saccharomyces cerevisiae* to express and deliver *E. tenella* antigens apical membrane antigen 1 (EtAMA1) [11], immune mapped protein 1 (EtIMP1) [12] and repeat 3 from microneme protein 3 (EtMIC3) [13] induced significant protection against high-level challenge in vaccinated Cobb500 broiler chickens [10]. However, the impact of vaccination and subsequent parasite challenge on the host gut and its enteric microbiota was not evaluated. Oral administration of heat-inactivated and freezed dried *S. cerevisiae* has previously been shown to ameliorate the effects of coccidiosis in broiler chickens while modulating the host immune response and microbiota [14, 15]. Understanding the influence of a yeast-vectored anticoccidial vaccine on host metabolome and microbiomes could therefore be used to inform future vaccine development.

Enteric microbiomes play crucial roles in shaping host physiological functions including provision of nutrients [16, 17], immune system maturation and regulation [18, 19]. *Eimeria* infection can cause imbalance in gastrointestinal ecosystems [20, 21], commonly referred to as dysbiosis, and raises the risk of enteric comorbidities such as necrotic enteritis caused by *Clostridium perfringens* [22]. Variation in the severity of damage caused by *Eimeria* infection has also been shown to associate with differences in enteric microbiomes. For example, high level caecal lesion scores recorded during *E. tenella* infection correlated with increased *Enterobacteriaceae* occurrence but decreased *Bacillales* and *Lactobacillales* [21]. However, little is known about physiological responses in gastrointestinal molecular and biochemical mechanisms, or variation in microbiota between immunologically naïve, infected and vaccinated chickens. Few studies have provided insight into chickens’ metabolic responses to infection or vaccination. Using an untargeted metabolomic profile assessment, Aggrey et al. (2019) found that carnitine-derived metabolites involved in fatty acid metabolism, and thromboxane B2, 12-HHTrE and itaconate involved in inflammatory responses, were influenced by *Eimeria acervulina* infection [23]. In the same way, a human shingles vaccine trial revealed that key metabolites such as sterol class metabolites, arachidonic acids, phosphoinositide, and diacylglycerol, were essential to immune signalling [24]. Here, we have created a multi-omics dataset defining caecal microbial populations (lumen contents and tissue-associated) and caecal tissue metabolomes using high-throughput sequencing of the 16S rRNA gene and liquid chromatography-mass spectrometry (LC-MS), respectively. We have used a Multi-Omics Factor Analysis (MOFA) [25, 26] machine learning model to systematically integrate data on caecal microbiota and the host metabolome sampled during an anticoccidial vaccine trial, investigating host microbe-associated signatures that can predict chicken health status and vaccine efficacy.

## RESULTS

### Caecal pathology and parasite load *post-Eimeria* challenge demonstrates efficacy of a candidate yeast-vectored anticoccidial vaccine

We previously evaluated the efficacy of an experimental *S. cerevisiae*-vectored anticoccidial vaccine using readouts of gut pathology (caecal lesion scores: 0-4), parasite replication (quantitative PCR of caecal tissue) and chicken performance (body weight gain, BWG) following oral challenge with 15,000 sporulated oocysts of *E. tenella* [10]. Briefly, lesion scores at 6 dpi were lower in vaccinated chickens compared to unvaccinated controls (V-C vs UV-C; p<0.001; Figure 1A). Parasite replication measured by qPCR as parasite genomes per host genome was also lower in vaccinated chickens at 6 dpi (p<0.001; Figure 1B). In contrast, BWG was not significantly different at 6 dpi (Figure 1C) although it was by 10 dpi [10].

**Figure 1.**
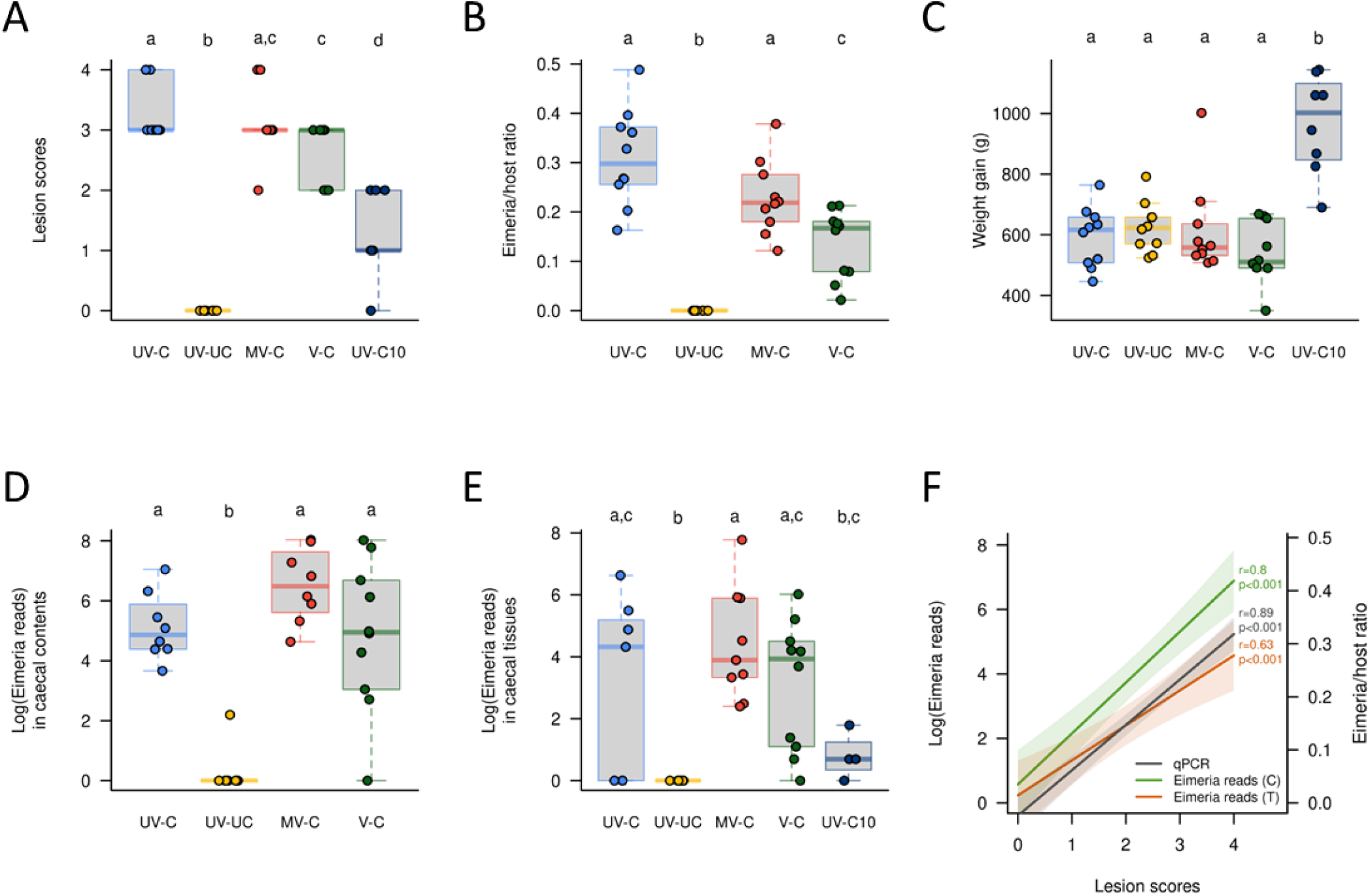
Summary of vaccine trial phenotypes assessed six days post infection (dpi) with 15,000 sporulated *Eimeria tenella* oocysts. (A) Caecal lesion scores, (B) parasite load represented as parasite genomes per host genome, determined using qPCR, (C) bodyweight gain from 0 to 6 dpi, (D and E) parasite load represented by *Eimeria* mitochondrial 16S rRNA sequence reads in caecal contents and tissue, (F) association between caecal lesion score and parasite load measures. Panels A-C reanalysed from Soutter et al., 2022. Groups UV-C: unvaccinated, challenged, UV-UC: unvaccinated, unchallenged, MV-C: mock vaccinated, challenged, V-C vaccinated, challenged.

In the present study, the level of *E. tenella* replication at 6 dpi was confirmed by quantification of *Eimeria* mitochondrial 16S rRNA amplicon reads in NGS microbiome data from caecal tissue and contents (Figures 1D and 1E). Comparison of all three *E. tenella* replication measures revealed a significant association with lesion score severity (qPCR ratio: r=0.89, NGS reads of caecal contents: r=0.8, NGS reads of caecal tissue: r=0.63; all p<0.001; Figure 1F). For comparison, 10 dpi unvaccinated and challenged chickens (UV-C10) considered to be recovering from infection also showed a significant reduction in gut pathology and *Eimeria* load compared to all infected subjects at 6 dpi (p<0.001; Figures 1A and 1E).

### Gut pathology and parasite load correlate with changes in gut microbiota

The composition of enteric microbial populations can reflect the health status of micro-ecosystems in the gastrointestinal (GI) tract. We performed 16S amplicon sequencing from caecal contents and tissues collected from the same individuals to characterize gut microbiota composition, with no significant differences in beta diversity detected between sample types (caecal tissue compared to caecal contents; PERMANOVA test R^2^=0.026, p=0.052) (Figure S1A). Comparison between caecal contents and tissue found 62.7-73.6% of microbiota composition to be shared (Figures S1B). Microbial populations enriched in caecal contents included *Lactobacillus mucosae*, *Lactobacillus salivarius*, *Paludicola psychrotolerans*, *Kineothrix alysoides*, *Anaerostipes butyraticus*, and [*Clostridium*] *polysaccharolyticum*; while microbial populations of *Anaerotruncus colihominis* (KTU 13) and *Flavonifractor plautii* (KTU 14) were enriched in caecal tissues (i.e. UV-C, MV-C, and V-C) (Figure S1C).

Principal coordinates analysis (PCoA) based on Bray-Curtis dissimilarity measurements showed that the caecal contents microbiota composition of unchallenged versus all challenged groups were distinct from each other (6 dpi) along the PCoA1 axis (31.15% of observed variation) (Figure 2). A PERMANOVA test confirmed significant differences in microbiota (R^2^=0.33, p=0.001) (Figure 2A) and there were significant correlations with caecal lesion scores (|r|=0.73, p<0.001), parasite load in caecal tissues (qPCR ratio: |r|=0.76, p<0.001) and caecal contents (NGS reads: |r|=0.67, p<0.001) (Figure 2B-D).

**Figure 2.**
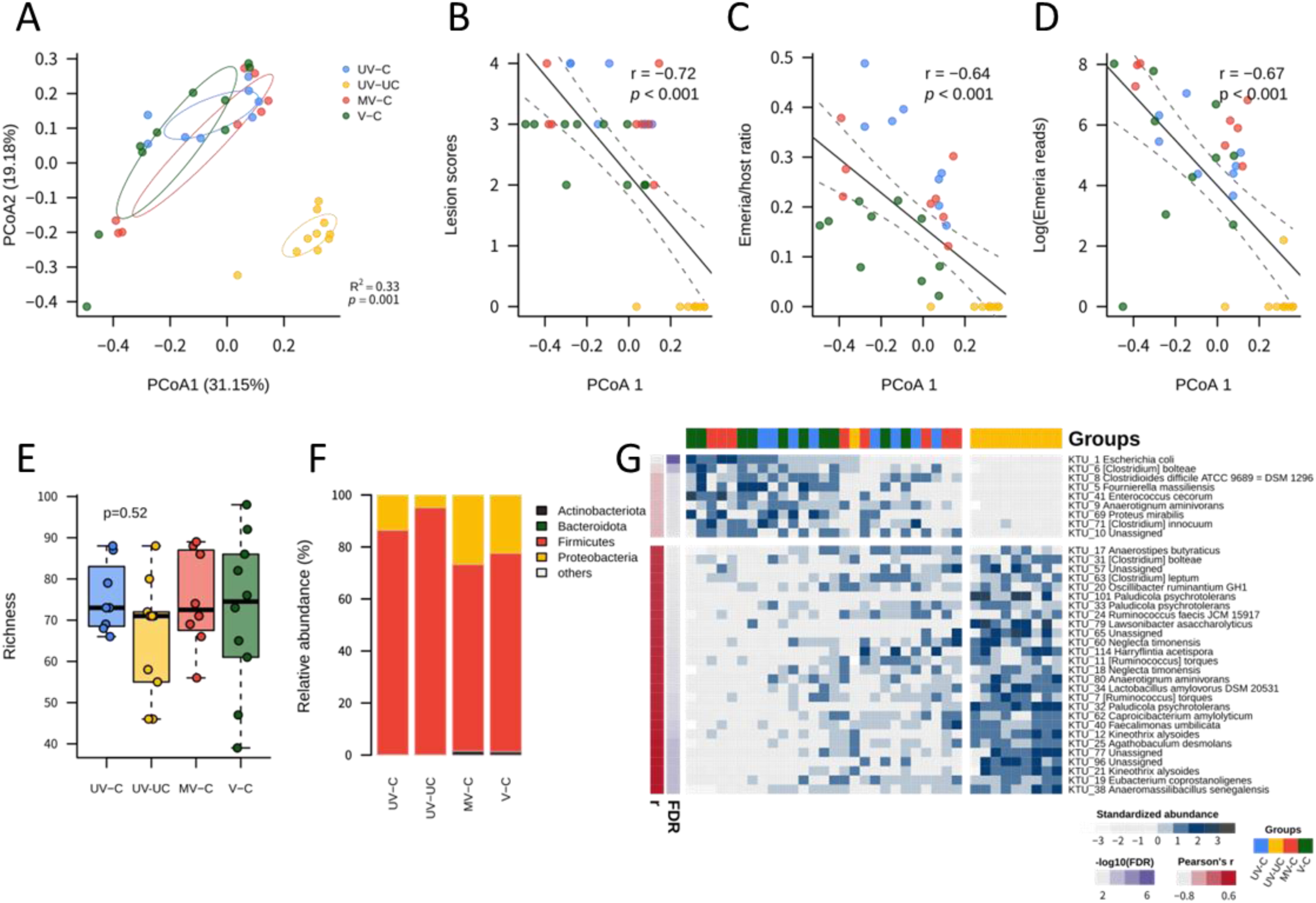
Gut microbiota profiling and associations of gut pathology and parasite loads. (A) Principal coordinates analysis (based on Bray-Curtis distance) for beta diversity of gut microbiota composition (caecal contents) among four groups of chickens. (B) The correlation between microbiota composition and lesion scores. (C)-(D) The correlations between microbiota composition and parasite loads, (C) based on qPCR quantification of the ratio of *Eimeria* and host genes, (D) based on NGS reads of *Eimeria* apicoplast 16S rRNA gene. (E) Alpha diversity (richness) of four groups of chickens. (F) Relative abundance of gut microbiota composition at the phylum level. (G) Gut pathology and parasite load associated microbes (Pearson’s r > 0.4 or < -0.4, FDR-adjusted p < 0.05) extracted from PCoA1 of panel (A). Groups UV-C: unvaccinated, challenged, UV-UC: unvaccinated, unchallenged, MV-C: mock vaccinated, challenged, V-C vaccinated, challenged.

On average a low alpha diversity index of microbial richness was found in all chickens across all groups (71.64±14.21) compared to a previous study by Hay et al. (2023) (493.13±201.60, re-analysed using the same pipeline used in the present study)[27]. This disparity may be due to requirement for broad-spectrum enrofloxacin treatment during this trial (Figure 2E). Comparison between the groups revealed a higher richness index in all challenged groups 6 dpi compared to the unvaccinated, unchallenged group (UV-UC), although the difference was not statistically significant. The dominant phyla were *Firmicutes*, followed by *Proteobacteria* in all chickens (combined, accounting for more than 98%) (Figure 2F); however, *Proteobacteria* were reduced in UV-UC chickens (4.83% comparing to 13.5%/26.65%/22.36% in other groups). *Actinobacteria* were enriched in both mock and true vaccinated groups (1.55% and 1.35%, respectively), dominated by genus *Bifidobacterium* (1.50% and 1.30%, respectively). Since the lesion scores and *Eimeria* loads were significantly correlated with the PCoA1 axis of beta diversity, 36 associated taxa enriched in challenged chickens were identified by Pearson’s correlation analysis (|r|≥ 0.4, FDR < 0.1), including *Escherichia coli*, *Clostridium difficile*, *C. innocuum*, and *Proteus mirabilis* (Figure 2G).

### Metabolomes reflect the molecular alterations of host physiology responses in health, infection, and recovery

Caecal tissue metabolomic profiling was performed for the same chickens as described above using samples collected in parallel with those used for microbiome sequencing analysis to characterize host physiological responses. An untargeted metabolomics approach was applied for screening metabolites within the tissues. Based on Euclidean distance measurements, PCoA of caecal tissue metabolome profiles showed a similar pattern to the caecal microbiota with unchallenged and challenged individuals differentiated along the PCoA1 axis (52.73% of observed variation) (Figure 3). The recovering (UV-C10) group displayed a broad but intermediate metabolome profile to that of 6 dpi challenged chickens and uninfected chickens and this group was also differentiated along the PCoA2 axis (9.57%). Host metabolome profiles correlated with caecal lesion scores (|r|=0.83, p<0.001) and *Eimeria* loads (qPCR ratio: |r|=0.73, p<0.001; NGS reads of caecal contents: |r|=0.83, p<0.001; NGS reads of caecal tissues: |r|=0.68, p<0.001) (Figure 3B-C). Among 1,180 metabolites belonging to the 10 categories that were detected from all chickens (including partially characterized and uncharacterized; Figure 3D), 954 metabolites were either negatively (606, non-infection-associated) or positively (348, infection-associated) correlated with pathophysiology changes (lesion scores and *Eimeria* loads; significant negative correlation with PCo1 in Figure 3A by Pearson’s correlation analysis, FDR < 0.1; Figure 3E). In more detail, xenobiotics, cofactors and vitamins, especially vitamin Bs, were characterized as non-infection-associated metabolites (Figure S2A and Table S2); while lipids, especially the sphingolipids, nucleotides, and carbohydrates, were characterized as infection-associated metabolites (Figure S2B and Table S2).

**Figure 3.**
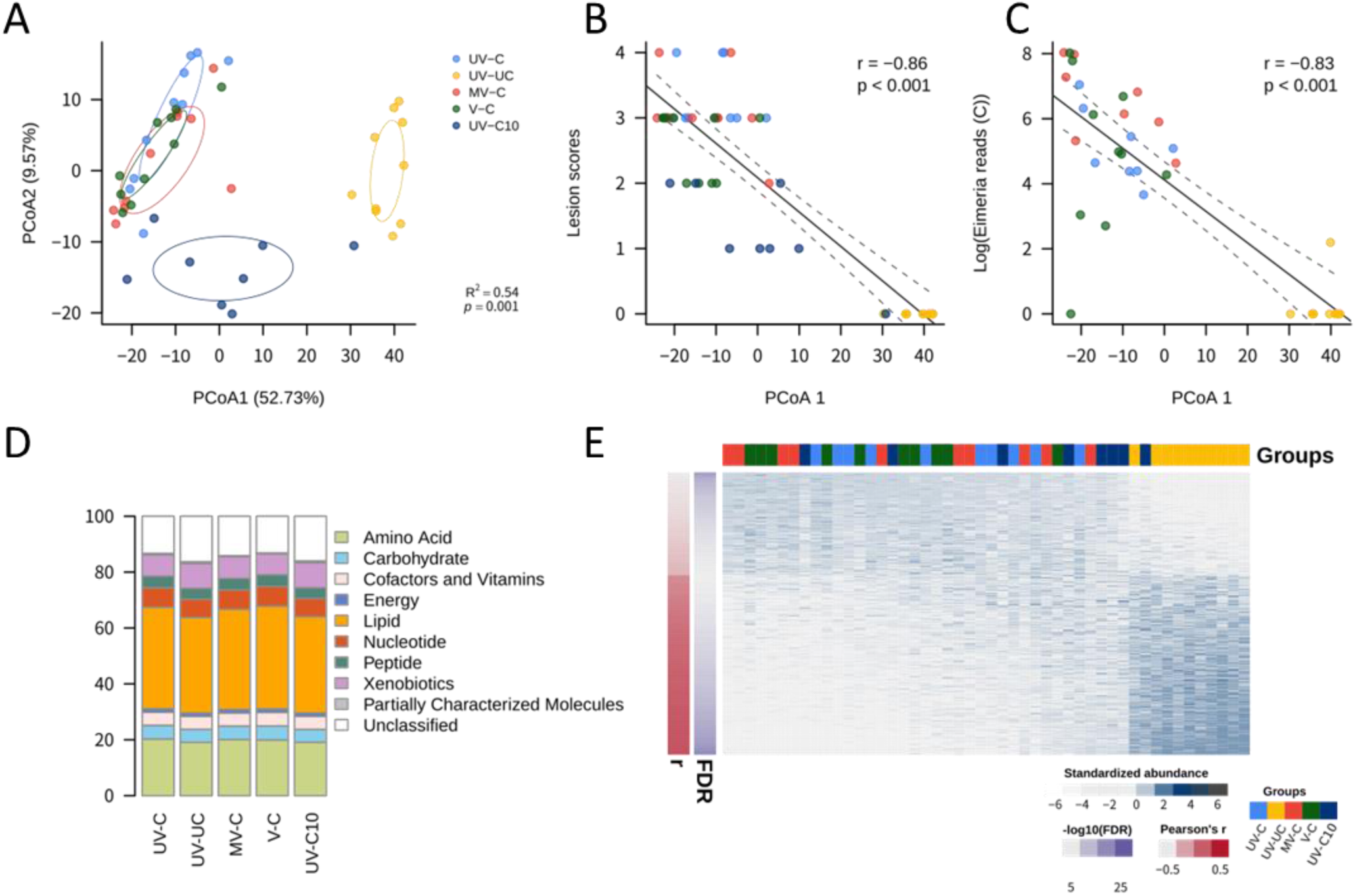
Chicken caeal tissue metabolome profiling and associations of gut pathology and parasite loads. (A) Principal coordinates analysis (based on Euclidean distance) for chicken caecal tissue metabolome composition among five groups of chickens. (B) The correlation between metabolome composition and lesion scores. (C) The correlation between metabolome composition and parasite loads, based on NGS reads of Eimeria apicoplast 16S rRNA gene. (D) Compositions and categories of the metabolome of five groups of chickens. (E) Gut pathology and parasite load associated metabolites (Pearson’s r > 0.4 or < -0.4, FDR-adjusted p < 0.05) extracted from PCoA1 of panel (A). Groups UV-C: unvaccinated, challenged, UV-UC: unvaccinated, unchallenged, MV-C: mock vaccinated, challenged, V-C vaccinated, challenged, UV-C10: unvaccinated, challenged, 10 days post-infection.

### Multi-omics factor analysis reveals covariation patterns of disease status

Using multi-omics factor analysis (MOFA), integration of parallel caecal tissue and content microbiomes with host tissue metabolome data showed concordant responses that associated with gut pathology and parasite load. Host-microbe intercorrelated features were assessed between microbial and metabolite features using Spearman’s correlation. A total of 151 KTUs and 767 metabolites were significantly associated (FDR<0.05), resulting in a MOFA model that contained 15 representative factors. The factors were decomposed and ordered by the fraction of significant associations they contributed to the major variances (Figure 4A). The first two MOFA factors explained the most variance that differentiated the unchallenged, challenged, and recovering groups on the MOFA scatter plot (Figure 4B). In addition, covariate (phenotype) correlation analysis demonstrated that the first two MOFA factors were associated with the majority of the covariates (Figure 4C) where factor 1 (FA1) was particularly associated with covariates related to infection (r < -0.6) and factor 2 (FA2) was associated with BWG (r=0.56); associations not identified in correlations of single omics analyses.

**Figure 4.**
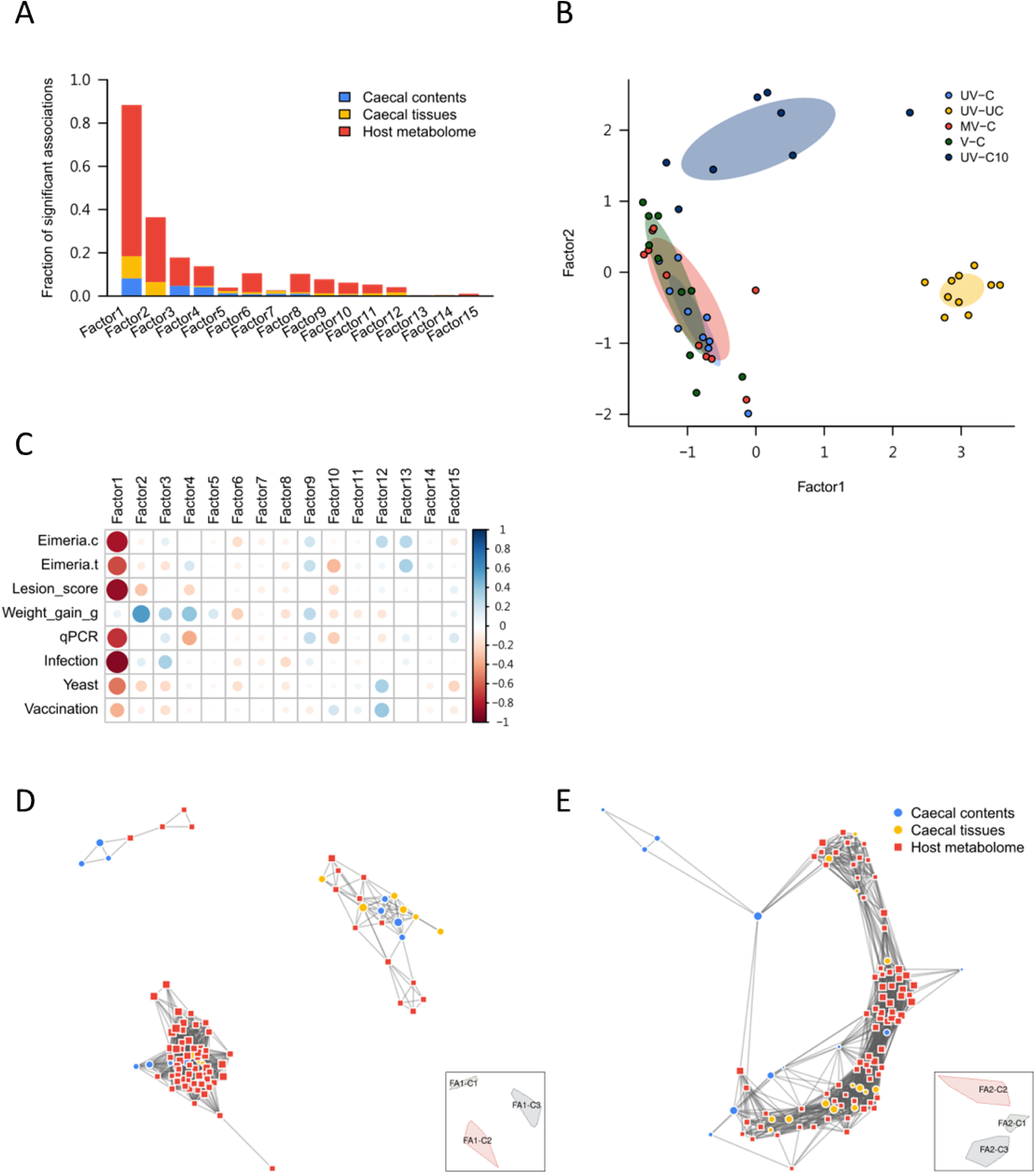
MOFA model for all trial groups and downstream signature marker identification by network analysis. (A) Bar plots showing the fraction of significant associations between the features of each microbiome or metabolome modality and each factor. The stacked bars interpret whether the variance-explained values are driven by a strong change in a small number of features or by a moderate effect across a large range of features. (B) Scatterplot of factor 1 (x axis) versus factor 2 (y axis). Each dot represents a sample, colored by the trial group. (C) The correlation heatmap of MOFA factors and phenotypes (Eimeria.c: NGS read-based *Eimeria* load in caecal contents; Eimeria.t: NGS read-based *Eimeria* load in caecal tissues; qPCR: qPCR-based *Eimeria* load in caecal tissues; Infection: infection condition-infected or non-infected; Yeast: yeast vector exposure or not; Vaccination: vaccination condition-vaccinated or non-vaccinated). (D)-(E) Network analysis and visualization for the features from (D) factor 1 and (E) factor 2. Thumbnail legends present the regions of subnetworks.

Multi-omics networks can contextualize the multiple types of microbiome disruption associated with various biological molecules found in different health statuses [28]. Additionally, a network’s hotspot molecular features (hubs and clusters) can highlight targets to be followed-up. Here, we conducted network analyses downstream of MOFA to explore biomarkers that might associate with anticoccidial vaccination. Network analyses for the MOFA factors showed sub-structures (clusters of intercorrelated features) that were enriched in each MOFA factor (Figure 4D-E). Three clusters were identified from FA1 components; two associated with *Eimeria* challenged chickens (including unvaccinated, vaccinated and recovering groups) (FA1-C1 and C3 in Figure 4D), whilst the third associated exclusively with unchallenged chickens (FA1-C2 in Figure 4D). Additionally, cluster 1 in the FA2 network demonstrated associations between unchallenged/recovering groups and the 6 dpi-challenged group (FA2-C1 in Figure 4E). Clustered components from the FA1 and FA2 networks associated with non-challenge and recovery were enriched by vitamin B and derivatives (e.g., pyridoxine, riboflavin, nicotinate derivatives), short-chain fatty acids (e.g., butyrate/isobutyrate and valerate), and short-chain fatty acid-producing bacteria (e.g., *Caproicibacter fermentans* and *Ruminococcoides bili*). Itaconate, an antipathogenic organic acid was enriched in recovering chickens. In contrast, uremic toxin (e.g., p-cresol sulfate), the long-chain fatty acids and derivatives (e.g., 14—18C fatty acids and glycerophospholipids (GPs), glycerophosphocholine (GPC), phosphoethanolamine (PE) derivatives), metabolites of fatty acid metabolism (eicosenoylcarnitine and docosadienoylcarnitine), and gut pathogens (e.g., *Clostridium difficile* and *C. innocuum*) and commensal bacteria (e.g, *Escherichia coli*, *Clostridium bolteae*, and *Fecalibacterium prausnitzii*), were enriched in post-*Eimeria* challenged associated clusters of both networks (Figure S3).

### MOFA models discover potential signature markers of host response to challenge after vaccination

While highlighting the covariation patterns of disease status, the MOFA model constructed using data from all samples did not reveal factors specifically associated with unvaccinated-challenged and vaccinated-challenged (mock and true vaccines) chickens. A more focused MOFA model was performed on all 6 dpi challenged groups to identify signature markers after vaccination. In the second model, the first four MOFA factors contributed to the major variation of the data and the fraction of significant associations (Figure 5A). Interestingly, the phenotypic and pathological covariates were more closely associated with FA4 and FA11 (e.g., lesion score severity was more associated with FA4 than other FA; r=-0.57). Vaccine treatment conditions (Yeast: treating with yeast vectors or not; Vaccination: treating with the true vaccine or not) were negatively associated with FA4 and FA11, and the parasite load (qPCR ratio) was associated with both FAs (r=-0.59 and -0.37) (Figure 5C).

**Figure 5.**
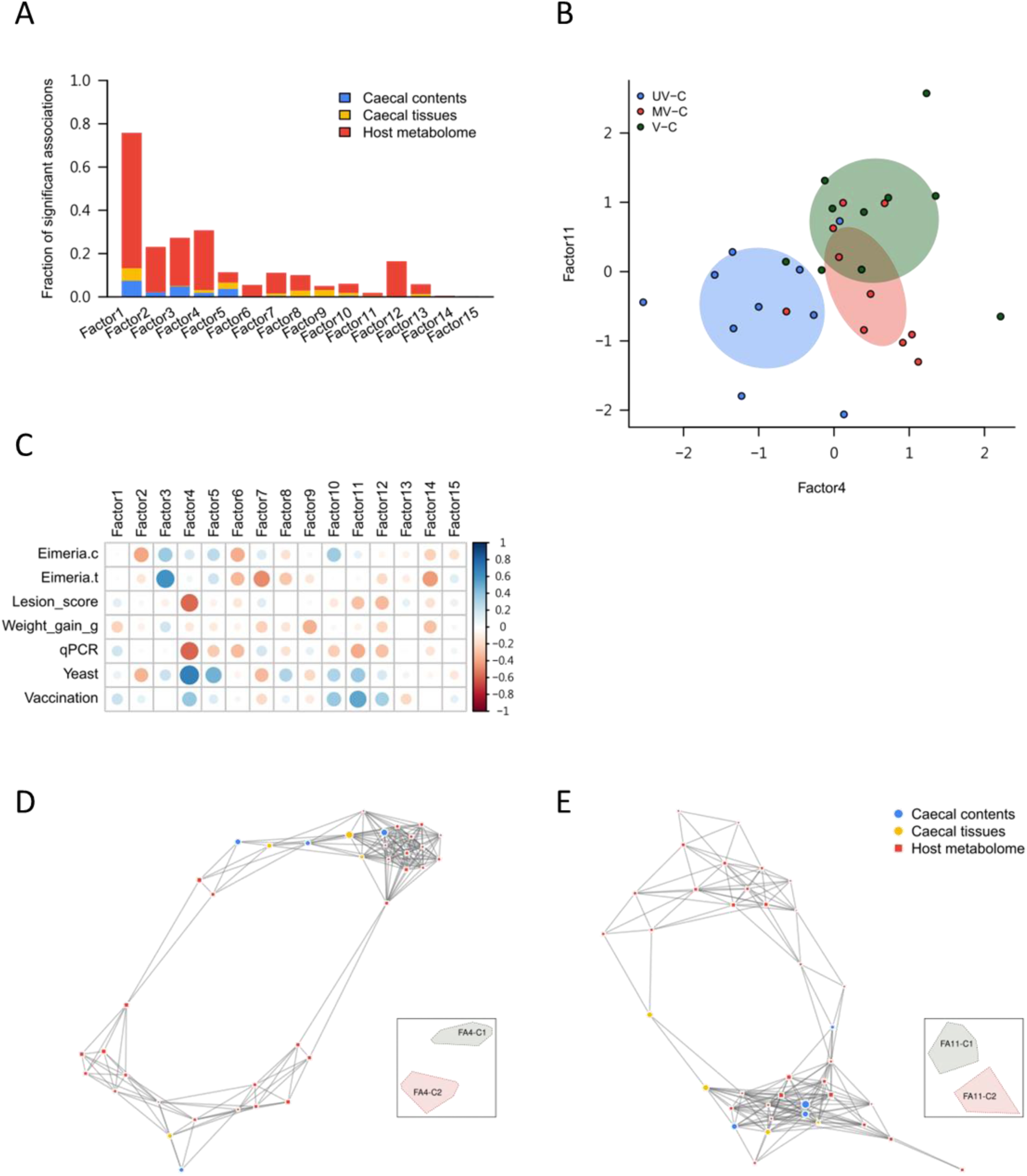
MOFA model for challenged 6dpi chickens and downstream signature marker identification by network analysis. (A) Bar plots showing the fraction of significant associations between the features of each microbiome or metabolome modality and each factor. The stacked bars interpret whether the variance-explained values are driven by a strong change in a small number of features or by a moderate effect across a large range of features. (B) Scatterplot of factor 4 (x axis) versus factor 11 (y axis). Each dot represents a sample, colored by the trial group. (C) The correlation heatmap of MOFA factors and phenotypes (Eimeria.c: NGS read-based *Eimeria* load in caecal contents; Eimeria.t: NGS read-based *Eimeria* load in caecal tissues; qPCR: qPCR-based *Eimeria* load in caecal tissues; Yeast: yeast vector exposure or not; Vaccination: vaccination condition-vaccinated or non-vaccinated). (D)-(E) Network analysis and visualization for the features from (D) factor 4 and (E) factor 11. Thumbnail legends present the regions of subnetworks.

Comparison of FA4 and FA11 using a scatter plot demonstrated that FA4 clearly distinguished the treatment condition of yeast vectors between unvaccinated (UV-C) and vaccinated groups (MV-C and V-C). FA11 showed a different trend between the mock vaccine group (MV-C) and the true vaccine group (V-C) (Figure 5B). Using network analysis, the signature features of various sphingolipids (e.g, sphingosine, sphingomyelin) and *Ruminococcus lactaris* were clustered from both FAs and enriched in most vaccinated subjects; whereas the long-chain fatty acids (e.g., linoleoyl-arachidonoyl-glycerol and oleoyl-oleoyl-glycerol) were enriched in unvaccinated-unchallenged chickens (Figure 5D-E, and Figure S4).

## DISCUSSION

An experimental yeast-vectored anticoccidial vaccine has recently been described as a step towards improved control of *Eimeria* species such as *E. tenella*, which cause coccidiosis in chickens [10]. Small-scale studies under commercial conditions found that vaccination could partially control the direct consequences of live parasite challenge, reducing parasite replication and its associated enteric pathology, while protecting performance (BWG and feed conversion ratio). In the present study, we have assessed the impact of vaccination on indirect consequences of *Eimeria* infection including microbial dysbiosis and metabolic disruption.

Using 16S rDNA amplicon sequencing from caecal contents and caecal tissue of experimentally vaccinated and challenged chickens (specific pathogen-free sourced), the microbiota richness was lower than in a previous farm study (Hay et al. 2023), likely due to the necessary enrofloxacin medication in the early rearing period to control an outbreak of colibacillosis. Indeed, it is well described that antibiotic treatment severly impacts on microbiome composition and richness, but recovering to its normal composition after stopping the treatment. While this was unexpected, such treatments are common under field conditions. Microbiota richness is also expected to be higher in populations of mixed breed chickens reared in the field under varied husbandry regimes than under the controlled conditions used in the present study. Comparison between caecal contents and tissues found no significant differences in alpha diversity (p=0.87) and beta diversity (p=0.052) (Figure S1). Only *Anaerotruncus (A.) colihominis* and *Flavonifractor (F.) plautii* were consistently enriched in caecal tissue samples across multiple groups (UV-UC, MV-C, and V-C), indicating their association with the intestinal mucosal environment. *A. colihominis*, originally isolated from mouse colonic mucosa by the Leibniz Institute DSMZ, has been detected in the intestinal lumen and stool samples of patients with bacteraemia and colorectal cancer, suggesting a potential broader role in gut dysbiosis and pathology [29, 30]. Similarly, *F. plautii*, known for its ability to degrade flavonoids and potentially mucins, was isolated by Levine *et al.* from mammalian intestinal mucosa [31]. Its presence in these tissue samples underscores its importance in gut health and disease [31, 32]. Based on our findings, investigation of caecal contents alone appears to be sufficient to investigate total gut microbiota because these reflect the primary condition of the intestinal ecosystem.

*Eimeria* infection is known to predispose chickens to diseases such as necrotic enteritis, caused by *C. perfringens* [33], and can disrupt enteric microbial populations leading to dysbiosis [21]. We anticipated that beta diversity, but not alpha diversity, would change following *Eimeria* challenge. However, although average richness was lower in unchallenged chickens, the difference was not statistically significant (Figure 2). Comparison of bacterial abundance between infected and non-infected chickens revealed increased Gammaproteobacteria and pathogenic *Clostridia* in *Eimeria*-challenged chickens. Common gastrointestinal pathogens, including *Escherichia (E.) coli*, *Clostridium (C.) difficile*, *Enterococcus (E.) cecorum*, *Proteus mirabilis*, and *Clostridium (C.) innocuum*, were also in higher abundance (Figure 2G) suggesting significant dysbiosis occurred following *Eimeria* challenge infection. It is notable that some strains of *E. caecorum* have been reported to cause high morbidity and mortality in broiler chickens [34]. Additionally, *C. difficile* and *C. innocuum* can cause antibiotic-associated diarrhea and have shown vancomycin resistance [35, 36], suggesting that a compromised gut environment may facilitate colonization by antibiotic-resistant strains; this condition mirrors the mechanism of human pseudomembranous colitis, which arises due to the overgrowth of *C. difficile* following extensive antibiotic usage. *Eimeria* infection can alter the gut microenvironment by increasing intestinal permeability and inflammation [37], thereby interacting bidirectionally with the gut microbiota. Consequences of enteric dysbiosis include immune dysregulation causing gut-related disorders such as allergies, inflammatory bowel disease (IBD) and autoimmune disorders [38–40]. Thus, *Eimeria* challenge is likely to activate a synergistic response between the host’s physiology and the commensal gut microbiota. Intestinal infections can decrease oxygen levels and lead to chronic tissue and mucosal hypoxia with dysregulation of activation of hypoxia-inducible factors (HIFs) and NF-κB, exacerbating inflammation and injury of intestinal tissues [41, 42]. The metabolic environment of the mucosa is also altered during inflammation since the Enterobacteriaceae require terminal electron acceptors from the mucosa for anaerobic respiration and blooming [43, 44]. Inflammatory cells release ROS and RNS, forming NO_3_^-^ as a terminal electron acceptor for Gammaproteobacteria growth via denitrification [43, 45–51].

In all *Eimeria*-challenged groups (UV-C, MV-C, V-C) the gut microbiota composition was similar (Figure 2A). However, yeast treatment groups (MV-C and V-C) showed a significant increase in *E. coli* abundance, nearly double in UV-C and over six times higher in UV-UC. While harmful *E. coli* may increase due to infection, it is possible that some protective *E. coli* strains that can stimulate an innate immune mechanism [52] and produce vitamins [53, 54] colonize after the reversion of dysbiosis. Notably, *Bifidobacterium*, a common lactic acid-producing probiotic, was present in both yeast treatment groups, irrespective of *Eimeria* antigen expression. In addition, Lactobacillales family bacteria (*Enterococcus*, *Lactobacillus*, *Pediococcus*) were enriched in both non-infected and yeast treatment groups, with *Lactobacillus* and *Pediococcus* being particularly higher in non-infected groups. This enrichment suggests a beneficial modulation of the gut microbiota. Yeasts and lactic acid-producing bacteria, often found together in nature [55], decrease pH value during fermentation creating an unfavourable environment for some pathogens [56, 57].

We used a multi-omics integrative tool, MOFA, to infer how the host metabolome interacts with gut microbes under a range of vaccination and *Eimeria* infection conditions. MOFA modelling confirmed that metabolites involved in fatty acid metabolism and β-oxidation pathways were altered by *Eimeria* infection [23]. Inflammation and oxidative stress induced by *Eimeria* invasion and subsequent pathology increase demand for metabolites involved in fatty acid metabolism [58]. The model found that carnitine derivatives such as eicosenolycarnitine and docosadienolycarnitine, intermediate metabolites involved in fatty acid metabolism, were enriched in the *Eimeria* challenged groups (challenge groups compared to non-challenge and recovering groups; factor 2 of MOFA model 1). In addition, p-cresol sulfate (pCS), a uremic toxin formed by gut microbial fermentation of tyrosine [59, 60], was also enriched in all challenged groups, especially in unvaccinated, challenged chickens (factor 1 of MOFA model 1). The main producer of pCS, *C*. *difficile*, a significant cause of diarrhoea during microbial ecosystem collapse, was also identified (factor 1 of MOFA model 1) [61, 62]. These findings link both layers of omics and prove evidence that *Eimeria* infection causes dysbiosis.

Since the first MOFA model (the full model with all groups of the trial) could not distinguish an effect of vaccination among the challenged, non-challenged and recovering groups, a second MOFA model was used to explore latent grouping among vaccinated and non-vaccinated chickens. We found sphingolipids, including sphingosine, sphingomyelin, and sphingoinositol, were significant factors associated with vaccination. Sphingolipids are required in cell membrane structures of eukaryotes (especially the Schwann’s cell, which surrounds the neuron axon) and some prokaryotes [63], as well as essential signalling molecules of inflammatory, immunity, cell autophagy, growth, and survival regulations [63–67]. Brown et al. (2019) indicated that the microbe-derived sphingolipids (especially from Bacteroides) are negatively correlated with gastrointestinal inflammation (i.e., inflammatory bowel disease) and maintaining homeostasis and symbiosis of gut microbiota [68]. This finding supports the efficacy of the yeast-based oral anti-coccidiosis vaccine and indicates that the vaccine can alter the symbiosis status of gut microbiota. However, only a few reads of Bacteroides were detected from yeast-based vaccine-treated samples and non-*Eimeria*-challenged samples (including from caecal tissues and contents), possibly due to the early antibiotic treatment of all study subjects. It implies that the microbial anti-inflammatory sphingolipids could be produced via other microbial species in the chicken gut microbiota, then act as a signal of anti-coccidiosis for the further applications.

In conclusion, using MOFA machine learning to integrate evaluation of potential interactions between the enteric microbiome and host metabolism provided a mechanistic insight into effects of anticoccidial vaccination and *Eimeria* challenge. In the present study, we identified Gamma-proteobacteria, *p*-cresol sulfate, *Bifidobacterium*, carnitine-derived metabolites and sphingolipids as host-microbe-associated biomarkers that vary between healthy, infected, vaccinated and/or recovering chickens, providing insights into potential strategies for controlling, treating and preventing coccidiosis. As we look to the future, the findings of this study are poised to contribute to the advancement of precision agriculture, particularly in enhancing poultry health management and the development of novel interventions against coccidiosis.

## MATERIALS AND METHODS

### Ethics statement

The animal experiments in this study were approved by the Royal Veterinary College (RVC) Animal Welfare Ethical Review Body (AWERB) and performed under the Animals in Scientific Procedures Act 1986 (ASPA) with a UK Home Office Licence.

### Study animals, metadata measurement and study design

Cobb500 broiler chickens were purchased from P. D. Hook (Hatcheries) Ltd. (Cote, UK) at day of hatch. All chickens received enrofloxacin (Baytril®, Bayer, Leverkusen, Germany, 10 mg Kg^-1^) from days 16 to 18 of the trial due to an outbreak of colibacillosis. Feeding and vaccination treatments were as described in a previous study (Study 4 in [10]). Briefly, four groups of ten chickens were sampled from a larger vaccination study six days post *E. tenella* challenge including (1) unvaccinated, challenged (UV-C), (2) unvaccinated, unchallenged (UV-UC), (3) mock vaccinated, challenged (MV-C), and (4) vaccinated, challenged (V-C) groups. A fifth group of eight unvaccinated, challenged chickens were sampled ten days post challenge (UV-C10; **Table S1**). Mock and experimental vaccines were administered by oral inoculation in 100 µl phosphate buffer saline (PBS) every 3-4 days from day 7 of age (five doses per chicken in total). Group 3 (MV-C) was vaccinated using a mock vaccine including *S. cerevisiae* EBY100 strain (Invitrogen, Thermofisher Scientific, Waltham, MA, USA) containing the empty yeast display plasmid vector pYD1 (Invitrogen). Group 4 (V-C) was vaccinated at the same timepoints by oral inoculation of an experimental trivalent formulation of *S. cerevisiae*-vectored recombinant vaccine using pYD1 to separately express each of three *E. tenella* antigens including EtAMA1 ectodomain [11], EtIMP1 [12] and EtMIC3 [13]. The vaccine design and administration procedures were as described previously [10]. Groups 1, 3, 4 and 5 were challenged by oral inoculation with 15,000 sporulated *E. tenella* Houghton strain oocysts at 21 days of age. Challenge oocysts were prepared and inoculated following established protocols [69]. Caeca (paired) were collected immediately post-mortem at six or ten days post infection (dpi, Groups 1-4, and 5, respecitively). The severity of infection was assessed using the Johnson and Reid scoring system [70]. Overall production performance was defined by Body Weight Gain (BWG) between 0 and 6 dpi. Parasite replication was measured using quantitative PCR for parasite genomes per host genome [10].

### DNA extraction and 16S amplicon sequencing

Bacterial genomic DNA was extracted separately from caecal tissue (∼100 mg) and caecal contents (∼200 mg) using a QIAamp Fast DNA Stool Mini kit (QIAGEN, Valencia, CA, USA) following the manufacturer’s pathogen detection protocol. 16S amplicon library preparation followed the Illumina 16S Metagenomic Sequencing Library Preparation guidelines [71]. The 16S ribosomal RNA (rRNA) gene V3–V4 hypervariable regions were amplified by PCR with the adapter overhang primers 341F (5′-TCGTCGGCAGCGTCAGATGTGTATAAGAGACAGC <u>CCTACGGGNGGCWGCAG</u>-3′) and 805R (5′-GTCTCGTGGGCTCGGAGATGTGTATAAGAGACAG <u>GACTACHVGGGTATCTAATCC</u>-3′) for 25 cycles. Indices and Illumina sequencing adapters were attached using the Nextera XT Index Kit with 8 cycles of a second amplification reaction. The final PCR products were purified using AMPure XP beads (Beckman Coulter, Brea, CA, USA). The amplicon DNA concentration was measured using Qubit dsDNA *HS* and *BR* Assay Kits (Thermo Fisher Scientific, Waltham, MA, USA). Library quality was determined using the Agilent Technologies 2100 Bioanalyzer system with a DNA-1000 chip. Eighty-eight samples representing caecal tissues from all chickens in Groups 1-5 (n=48) and caecal contents from all chickens in Groups 1-4 (n=40) were pooled with equal molality. The 16S amplicon libraries were sequenced using a 301 bp paired-end (301bp x 2) approach on an Illumina MiSeq platform using V3 chemistry.

### Bioinformatic processing and microbiota analyses

The Illumina MiSeq platform generated a total of 22,525,182 paired-end sequences. Sequences were cleaned by sequence length ≥ 300bp using Trimmomatic [72]. The 16S amplicon sequences were processed using the Quantitative Insights Into Microbial Ecology 2 (QIIME 2) pipeline (version 2019.10) [73]. Primer sequences were removed by Cutadapt (version 1.15) [74]. Trimmed sequences were truncated at 240 bp (forward) and 210 (reverse) and denoised using the DADA2 algorithm [75]. Amplicon sequence variants (ASVs) were obtained via the denoising process with quality filtering and chimera removal. A k-mer based re-clustering algorithm ‘KTU’ [76] was subsequently applied to assemble ASVs into optimal biological taxonomic units (KTUs). KTUs taxonomy was assigned by comparison with the SILVA SSU reference nr99 (v138) [77, 78] and NCBI 16S RefSeq (retrieved 10^th^ Feb. 2022) databases using the kaxonomy function of the KTU R-package. Eukaryotic organelle 16S sequences (identified as *Eimeria*) were extracted and used for supplementary parasite load quantification; non-prokaryotic and unassigned KTUs were removed from the microbiota dataset. The 309 KTU microbiota dataset was rarefied at the minimum read counts among samples (10,034 reads) after removing twelve samples with shallow sequence depth (< 10,000 reads).

Microbiota analyses were conducted and visualized using the Microbiome Analysis R code (MARco) [79], Community Ecology ‘vegan’ [80], and Pretty Heatmap (pheatmap) [81] packages in R (version 4.0.1) [82]. The ANOVA test with Tukey HSD *post-hoc* multiple comparison test or Kruskal-Wallis test with Dunn’s *post-hoc* multiple comparison test were used for parametric and non-parametric statistical analyses of group comparisons with a significance level of α = 0.05, and the P values were adjusted with a false discovery rate (FDR). Alpha diversity indices were estimated by richness. Beta diversity of microbial communities was measured by Bray-Curtis dissimilarity using principal coordinates analysis (PCoA), and heterogeneity was tested using ADONIS and ANOSIM tests.

### Metabolome profiling

Untargeted metabolome profiling of caecal tissues was performed by Metabolon (NC, USA) using their vendor protocol. Briefly, all samples were deproteinized by dissociating small molecules bound to protein or trapped in the precipitated protein matrix. To recover chemically diverse metabolites, methanol was used for protein precipitation under vigorous shaking for 2 min (Glen Mills GenoGrinder 2000) followed by centrifugation. The extract was aliquoted into five fractions: two for analysis by separate reverse phase (RP)/UPLC-MS/MS methods with positive ion mode electrospray ionization (ESI), one for analysis by RP/UPLC-MS/MS with negative ion mode ESI, one for analysis by HILIC/UPLC-MS/MS with negative ion mode ESI, and one sample was reserved as backup. Samples were placed briefly on a TurboVap® (Zymark) to remove the organic solvent. The sample extracts were stored overnight under nitrogen before preparation for analysis.

All methods used Waters ACQUITY ultra-performance liquid chromatography (UPLC) and a Thermo Scientific Q-Exactive high resolution/accurate mass spectrometer interfaced with a heated electrospray ionization (HESI-II) source and Orbitrap mass analyzer operated at 35,000 mass resolution. Each sample extract was dried then reconstituted in solvents compatible to each of the four methods. Each reconstitution solvent contained a series of standards at fixed concentrations to ensure injection and chromatographic consistency. One aliquot was analyzed using acidic positive ion conditions, chromatographically optimized for more hydrophilic compounds. The extract was gradient eluted from a C18 column (Waters UPLC BEH C18-2.1x100 mm, 1.7 µm) using water and methanol, containing 0.05% perfluoropentanoic acid (PFPA) and 0.1% formic acid (FA). Another aliquot was also analyzed using acidic positive ion conditions; however, it was chromatographically optimized for more hydrophobic compounds. The extract was gradient eluted from the same afore mentioned C18 column using methanol, acetonitrile, water, 0.05% PFPA and 0.01% FA and was operated at an overall higher organic content. Another aliquot was analyzed using basic negative ion optimized conditions using a separate dedicated C18 column. The basic extracts were gradient eluted from the column using methanol and water, however with 6.5mM ammonium bicarbonate at pH 8. The fourth aliquot was analyzed via negative ionization following elution from a HILIC column (Waters UPLC BEH Amide 2.1x150 mm, 1.7 µm) using a gradient consisting of water and acetonitrile with 10mM ammonium formate, pH 10.8. The MS analysis alternated between MS and data-dependent MS^n^ scans using dynamic exclusion. The scan range varied slightly between methods, but covered 70-1000 m/z.

Raw data were extracted, peak-identified and QC processed by Metabolon’s in-house systems. Compounds were identified by comparison to library entries of purified standards or recurrent unknown entities. The in-house library was built and maintained by Metabolon, and contained more than 3,300 commercially available purified standard compounds with the information of retention time/index (RI), mass to charge ratio (*m/z)*, and chromatographic data (including MS/MS spectral data). Compound identification was based on the following criteria: retention index within a narrow RI window of the proposed identification, accurate mass match to the library +/-10 ppm, and the MS/MS forward and reverse scores between the experimental data and authentic standards.

A subset of 1,180 metabolites was detected from the untargeted metabolomics screen. Each metabolite’s peak area (i.e. total ion counts, integrated area-under-the-curve) was median-scaled to normalize. The missing values were then imputed with the observed minimum of each metabolite. Since the metabolomic data were typically close to log-normal distribution, the normalized-imputed data were transformed using the natural log for subsequent analyses.

### Multi-Omics Factor Analysis (MOFA) model for microbiota and metabolome integrative analysis

MOFA model fittings were performed to integrate multi-omics data modalities based on an unsupervised machine learning model formulated in a probabilistic Bayesian framework. The 16S rRNA amplicons of caecal tissue and content, and host caecal metabolome were the separate data modalities in this study. In order to make all omics data comparable, the amplicon abundance was centered log-ratio transformed using the ‘clr’ function of the compositions R-package. Spearman’s correlation (FDR <0.05) was implemented to select associated features from the omics datasets [83]. Downstream characterization was performed by variance decomposition, detecting the fraction of significant associations between the features and each factor using Pearson’s correlation (FDR < 0.1), and correlation of phenotype covariates. A sub-grouped MOFA model fitting was performed on all 6 dpi challenged groups. A network analysis for identifying sub-structures of MOFA factors was performed with the R package igraph47 [84]. An adjacency matrix based on Spearman’s correlation coefficients of intercorrelated features was constructed from a MOFA factor of interest; these coefficients were also used for assessing length of edges on the network. The latter was conducted with the fast greedy modularity optimization algorithm [85] to identify clusters in the network.

## SUPPORTING INFORMATION

**Figure S1.**
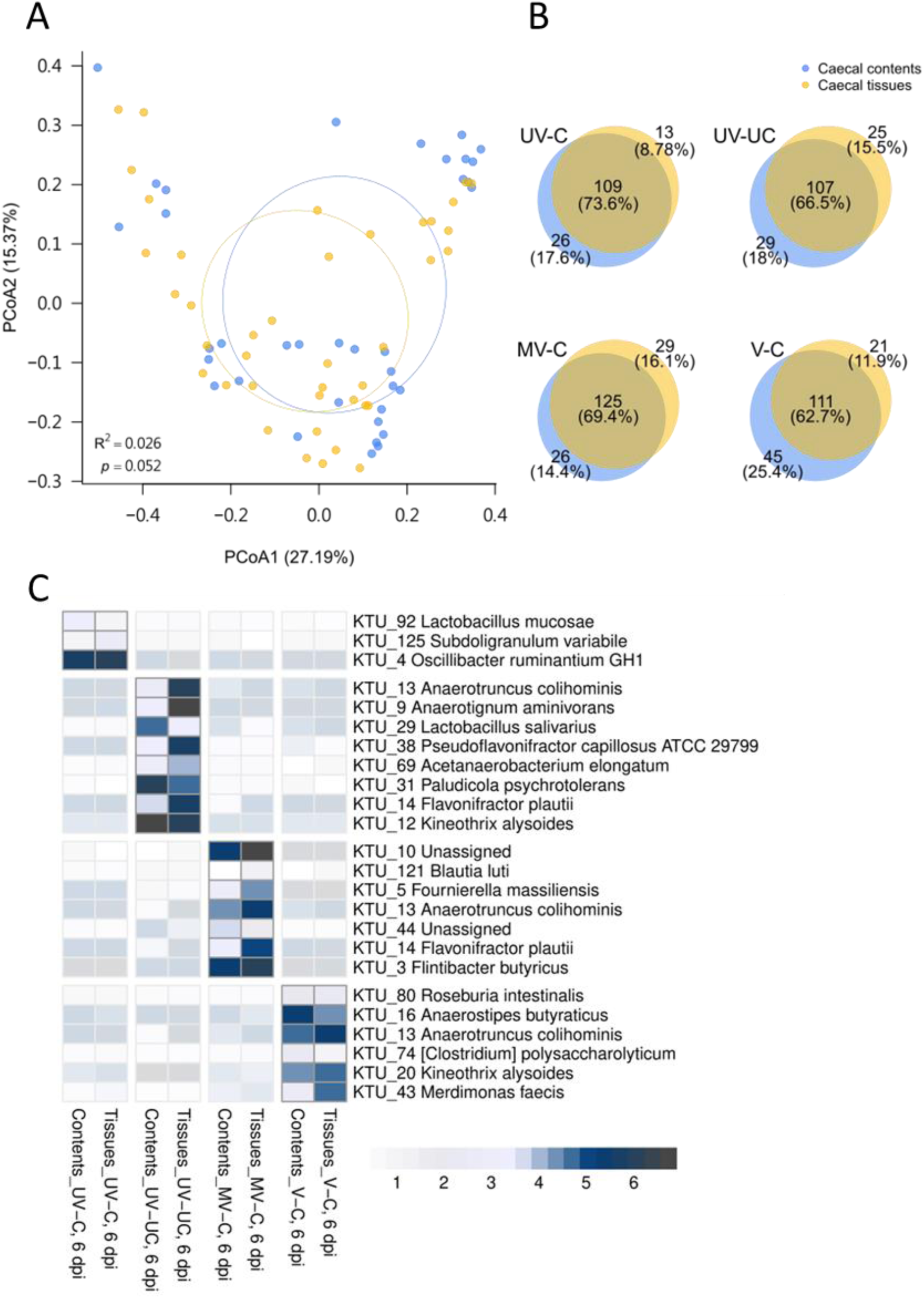
Comparisons of microbiota profiles between chicken caecal contents and caecal tissues. (A) Beta diversity analysis with the PERMANOVA test demonstrateed no significant differences between caecal contents and caecal tissues (p=0.052). (B) Venn diagrams show 62.7-73.6% microbiota composition shared between caecal contents and tissues in four study groups (UV-C, UV-UC, MV-C, and V-C). (C) Differential abundance (p<0.05 by DESeq2 test) of microbes between caecal contents and caecal tissues in four study groups, respectively.

**Figure S2.**
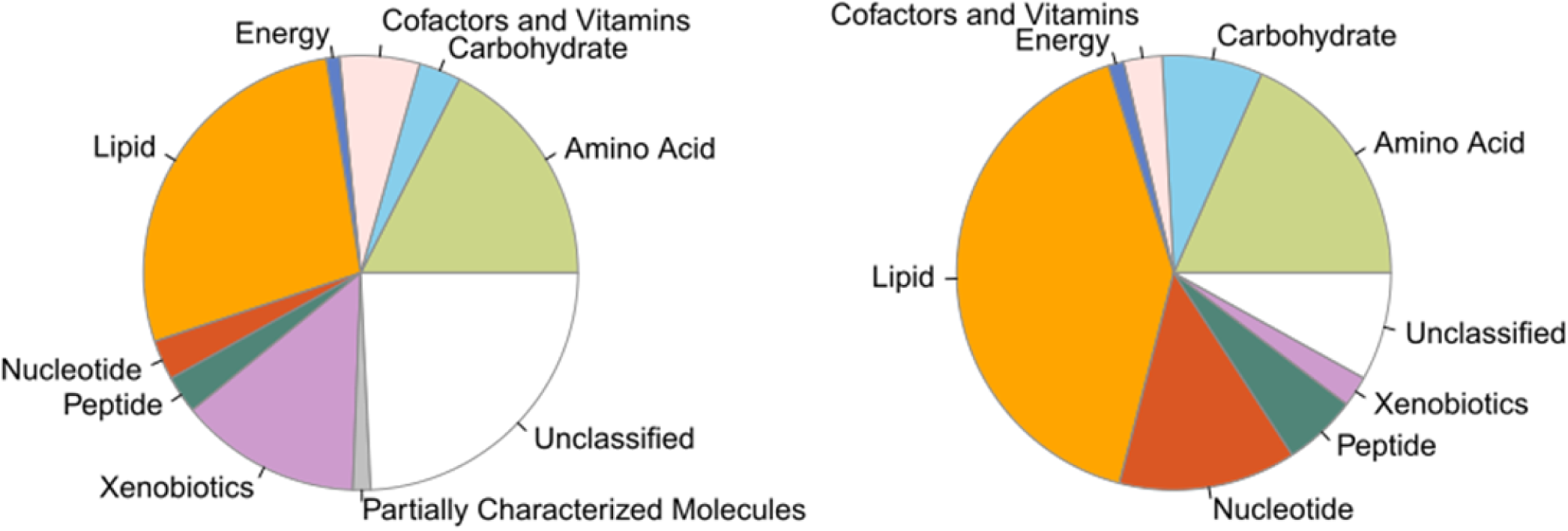
Lesion score and *Eimeria* load correlated metabolites. (A) Categories of non-infection-associated metabolites. (B) Categories of infection-associated metabolites.

**Figure S3.**
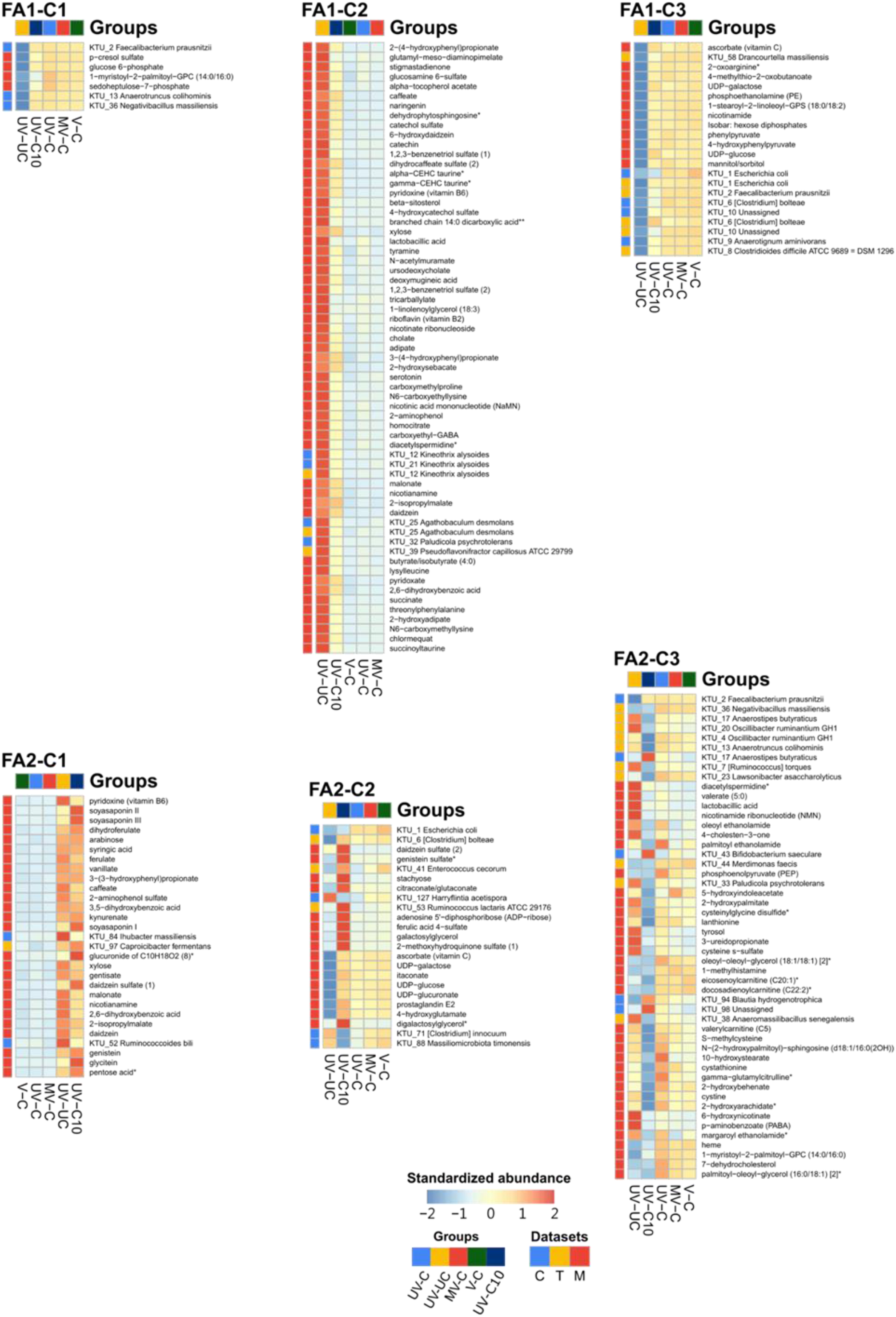
Details of subnetworks from the MOFA model. **Microbial and** metabolomic features of each cluster (C1-C3) in FA1(Figure 4D) and FA2(Figure 4E)— the abundance of microbial and metabolomic features among groups were presented by heatmap. The left columns were annotated by sample sources (C: caecal content microbiome; T: caecal tissue microbiome; M: caecal tissue metabolite).

**Figure S4.**
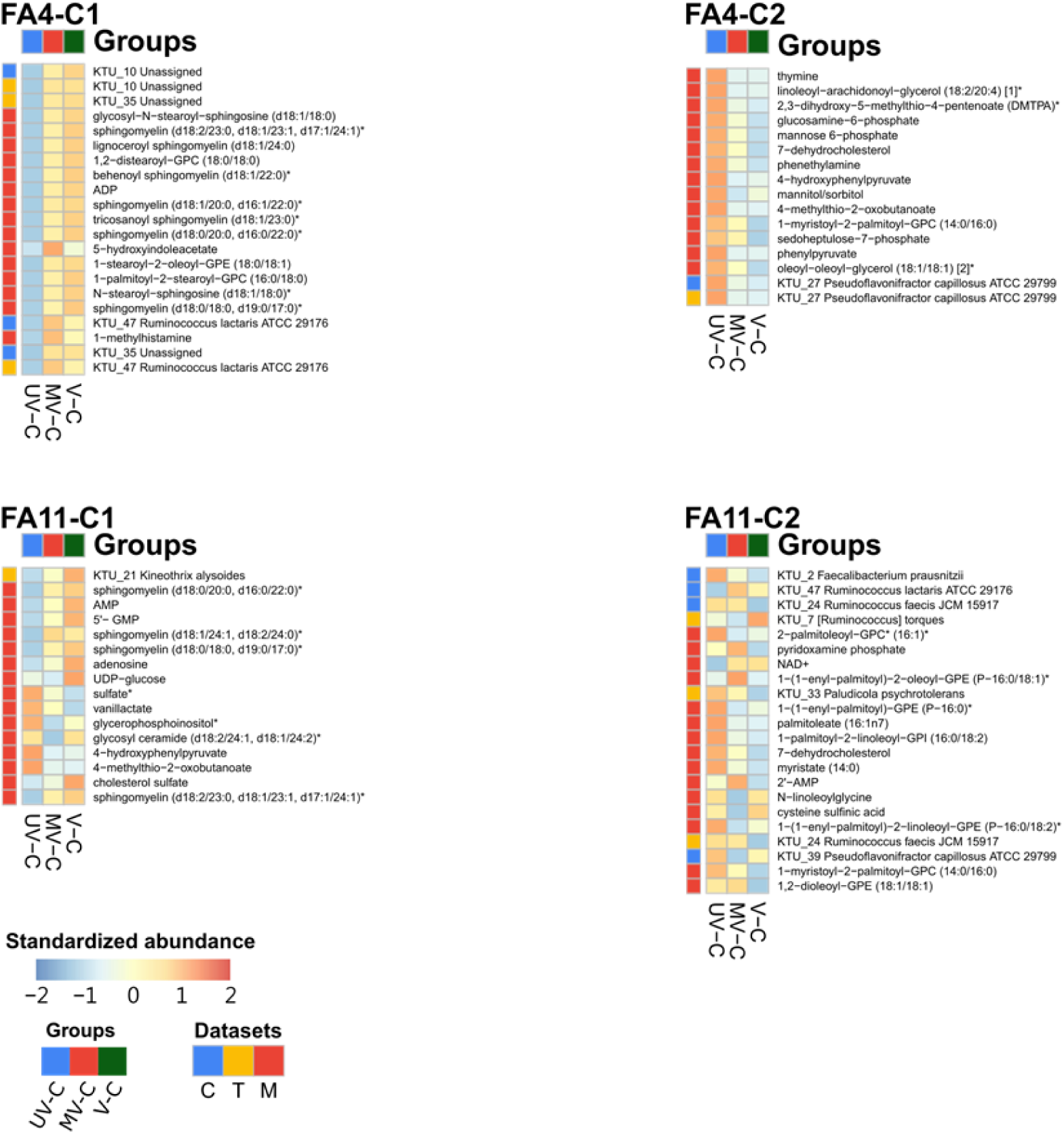
Details of subnetworks from 6dpi -challenged MOFA model. Microbial and metabolomic features of each cluster (C1-C2) in FA4(Figure 5D) and FA11(Figure 5E)—the abundance of microbial and metabolomic features among groups were presented by heatmap. The left columns were annotated by sample sources (C: caecal content microbiome; T: caecal tissue microbiome; M: caecal tissue metabolite).

**Table S1.** Sample information of studying groups.

| Groups | # of samples | Description | Content ID | Tissue ID |
| --- | --- | --- | --- | --- |
| UV-C | 10 | Unvaccinated, challenged, 6 dpi | C1 | T1 |
| UV-UC | 10 | Unvaccinated, unchallenged, 6 dpi | C2 | T2 |
| MV-C | 10 | Mock vaccinated, challenged, 6 dpi | C3 | T3 |
| V-C | 10 | Vaccinated, challenged, 6 dpi | C4 | T4 |
| UV-C10 | 8 | Unvaccinated, challenged, 10 dpi | - | T5 |

**Table S2. Infection-associated metabolites**

## DATA AVAILABILITY

The original data set presented in the study is publicly available. These data can be found at NCBI under BioProject accession number: PRJNA990995.

## ACKNOWLEDGEMENTS

We thank Dr Meiyeh Jade Lu from the NGS Core, Academia Sinica, and Ms Yu-Tang Yang from the National Taiwan University College of Medicine for the technical consultancy of the NGS library construction. We gratefully acknowledge the support of the Houghton Trust Research Grants for funding this project.

## AUTHORS’ CONTRIBUTION

Conceptualization: P-Y.L., D.X., F.M.T., and D.P.B. Performed experiment: P-Y.L., J.L., F.S., J.J.O, and D.P.B. Data analysis: P.-Y.L. Contributed data or analysis tools: F.S., D.W., and, O.G. Writing—original draft: P-Y.L. and D.X. Writing—review and editing: P-Y.L., D.X., and D.P.B.

